# Introduction of a precocious metamorphosis BmNPV infection in the latter half of the fifth instar silkworm larvae

**DOI:** 10.1101/2020.09.24.311290

**Authors:** Pingzhen Xu, Meirong Zhang, Xueyang Wang, Yangchun Wu

## Abstract

The silkworm, *Bombyx mori*, is a complete metamorphosis insect, the model to study insect physiology and biochemistry. *Bombyx mori* nucleopolyhedrovirus (BmNPV) is a principal pathogen of the silkworm and its host range is restricted to silkworm larvae, requiring interaction with larvae to accomplish virus replication. Prothoracic glands (PGs) are a model for synthetic ecdysone with regulating insect growth and development. This study performed a transcriptome analysis of silkworm PGs after BmNPV infection. Transcriptome data were annotated with KEGG, GO, and shown to be of high quality by RT-qPCR. The spatial expression profiles of *BmJing* and *BmAryl* indicate that they may be specifically expressed in silkworm PGs. The RT-qPCR results of the DEGs in the PGs of BmNPV-infected larvae at 24, 48, and 72 h and at the developmental stages of days-6 and 7, comparing to day-3, reveal that the DEGs may be related to the BmNPV infection via promoting early maturation in the latter half of the silkworm fifth instar. This study is the first report on the identification of possible genes in PGs correlating with the precocious molting and metamorphosis of silkworm larvae under BmNPV infection in the latter half of the fifth instar. Our findings will help to address the interactions between BmNPV infection and host developmental response. This work provides a new perspective on BmNPV infection and host developmental response, as well as suggesting candidate genes for further research.

## Introduction

In insects, dietary sterols including cholesterol and phytosterols are essential substrates for insect ecdysteroids synthesis in prothoracic glands (PGs), which are important endocrine organs [1]. There are variations in the exact molecule of ecdysteroids that are secreted by PGs [2]. Ecdysteroids play various physiological roles in regulating growth, moulting, and metamorphosis [1, 3]. In *Drosophila melanogaster*, the corpora cardiaca (CC), corpora allata (CA) and prothoracic gland (PG) are fused into a complex endocrine structure known as a ring gland (DmRGs) [4]. In the domesticated silkworm *Bombyx mori*, one of the model species of insects, the PGs are a pair of semi-transparent or transparent saccate cell clusters that are located in the tracheal clusters of the prothorax.

The PGs known to be cholesterol-rich tissue are the main site of synthetic ecdysteroids from cholesterol and their subsequent secretion [5, 6]. On account of ecdysteroids synthesized from cholesterol, the supply of cholesterol to a steroidogenic organ is a major step for ecdysteroids production [1]. Cholesterol is a main sterol in insects. Insects lack the ability to synthesize their own sterols [7]. In lepidopteran, e.g., phytophagous *Manduca sexta* and *B. mori*, cholesterol is generated via the dealkylation of plant sterols in the gut and thereby utilized in ecdysteroid synthesis [7]. In the silkworm, dietary sterols are converted to cholesterol in the midgut and are the initial source of dietary sterols for other tissues [8, 9]. Cholesterol is principally detected in the sterol compositions of major lipid-related tissuses such as the midgut, hemolymph, and fat body of silkworm larvae during the feeding period [9]. The tissues of silkworms have tissue-intrinsic sterol profiles [9].

Cholesterol is rarely metabolized in the silkworm body and is primarily used for conversion to ecdysteroids [10]. An ecdysteroid (a steroid hormone) is required for molting and metamorphosis, and this hormone is synthesized in PGs [9]. Steroidogenesis has been well demonstrated for cholesterol biotransformation into ecdysone in silkworm larvae PGs [1]. Ecdysteroid is synthesized from cholesterol through a multistep conversion and most of its intermediates have been clarified. However, in silkworm PGs, only 7-dehydrocholesterol (7dC) is detected among the intermediates of ecdysteroid [11]. Cholesterol is converted into 7dC at the first enzymatic reaction with a single enzymatic reaction, and then finally transformed into ecdysone, which is a prohormone of ecdysteroid in the PGs [9]. In silkworm PGs, the primary features are the appearance of 7dC and the high level of cholesterol, and the PGs tissues show a high preference for cholesterol and 7dC [9]. This may be linked to the status of the main site of synthetic ecdysteroids, and PGs might possess a management system allowing for high retention of cholesterol and 7dC [9, 12]. The prohormone of ecdysteroids produced in PGs is secreted into the hemolymph and converted into the active form of 20-hydroxyecdysone (20E) [9].

Moreover, cholesterol is a necessary component of cell membranes [13], and it is also regarded as a constituent of active regions for host–virus interactions and increases the infectivity of viruses during the entry of enveloped viruses [14–16]. Both the cellular cholesterol level and cholesterol are identified as key factors for the *Bombyx mori* nucleopolyhedrovirus (BmNPV) when invading host silkworm larvae [17–19].

BmNPV is an enveloped double-stranded DNA virus and is the pathogen of a disease called silkworm grasserie, a disease which has led to severe damage to the sericulture industry in China [20, 21]. BmNPV produces two distinct forms in its lifecycle, namely, the occlusion derived virus (ODV) and budded virus (BV) [22]. Both forms have different roles during pathogenesis. The ODV is packaged in viral polyhedral bodies and is responsible for primary infection through the oral transmission of the virus among silkworm larvae [22], while the BV includes one or more nucleocapsids in a single envelope and is responsible for the secondary infection, causing systemic spreading all over the host within the infected silkworm larvae [22].

BmNPV is a principal pathogen of the silkworm and its host range is restricted to silkworm larvae [22]. BmNPV requires interaction with silkworm larvae to accomplish virus replication. The previous studies investigating the interactions between BmNPV and its host have mainly focuses on the first half of the fifth instar silkworm larvae. The PGs are a model for synthetic ecdysone for regulating insect growth and development. Several studies have made transcriptome analysis results available for silkworm PGs of model insect species, taking advantage of prior developments [23–26]. In the present study, we firstly investigate the precocious molting and metamorphosis of silkworm larvae under BmNPV infection in the latter half of the fifth instar and then use RNA sequencing (RNA-seq) to analyze silkworm PGs. This analysis is carried out in order to understand the complex biological processes in the interactions between BmNPV and its precocious metamorphic hosts.

## Materials and Methods

### Insects and virus

*B. mori* F_50_ strain (yellow blood) larvae were reared on fresh mulberry leaves under a 12:12 L: D photoperiod at 25 ± 1 °C and 60% relative humidity. The majority of the fifth instar larvae started wandering on day 7, depending on the batch of the silkworm. The larvae underwent oral inoculation with a wild BmNPV T3 strain and the occlusion-derived virus (ODV) was obtained from the larvae hemolymph before the larvae died. The ODVs were purified by repeated and differential centrifugation, as previously described [22].

### Collection of samples

The head, integument, midgut, fat body, hemocyte, ovary, testis, malpighian tubule, trachea, ASG (anterior silk gland), MSG (median silk gland), and PSG (posterior silk gland) of day-3 fifth instar larvae were collected for tissue expression analysis of differentially expressed genes (DEGs). Overall, 500 of the day-4 fifth instar larvae were orally infected with 200 ODVs of the BmNPV T3 strain per larva. The 500 control larvae were fed with the same volume of sterile distilled water instead of the ODV. The two groups were kept in isolation and reared under the same condition. The PGs were entwined in the tracheal bush of the first spiracle in pairs (**S1 Fig**). The PGs were carefully removed from the larvae of the infected and control groups after 24, 48, and 72 h (**S2 Fig**). The PGs were collected from the day-3 (V3), day-6 (V6), and day-7 (V7) fifth instar larvae for different developmental stage expression analysis of differentially expressed genes.

### Statistics of precocious maturation of silkworm after infection

Day-4 fifth instar larvae were divided into six groups with 200 in each group. All 200 larvae used in each of the three independent experiments were orally infected with 200 ODVs of the BmNPV T3 strain per larva. The 200 control larvae used in each of the three independent experiments were fed with the same volume of sterile distilled water instead of the ODV. The infected and control groups were kept in isolation and reared under the same conditions. The diseased and dead silkworms were removed and counted during rearing. When the proportion of mature silkworms was greater than 5% (first gate), the statistics started and the timing was set to 0 hours. Then, the number of the majority of larvae that had started maturing (second gate) was also counted. In the experiment for investigating the mature changes of the fifth instar larvae, only the numbers of the first gate matured silkworms and second gate matured silkworms were used.

### Cholesterol and 7-dehydrocholesterol feeding experiments

According to that previously described [27], mulberry leaves supplemented with 8000 mg/L of cholesterol and 7-dehydrocholesterol (7dC) were fed to the silkworm larvae, respectively. Mulberry leaves supplemented with the same volume of sterile distilled water were used as a control. Day-5 fifth instar larvae were fed with cholesterol and 7dC for the first time and then fed for a second time 24 hours later. Replacement mulberry leaves were added 6 hours after each feeding. Then, the number of the majority of larvae that had started maturing was counted. For each group, including the 200 larvae, the experiment was repeated three times.

### Examination of viral DNA in PGs

Reverse transcription-PCR (RT-PCR) was used to analyze the BmNPV virus replication level. Total DNA was extracted from the PGs of the BmNPV-infected *B. mori* larvae at 24, 48, and 72 h, as well as from the control larvae at 48 h. The DNA templates (10 ng) were PCR-amplified using primers for the *BmNPV IE1*, *GP41*, and *GP64* genes. The silkworm *glyceraldehyde-3-phosphate dehydrogenase* (*BmGAPDH*) was used as an internal control. The specific primers for each gene used in the RT-PCR are shown in **S1 Table**. The RT-PCR product of each gene was defined as previously described [28].

### Transcriptome analysis

Total RNA was isolated from the PGs of the silkworm larvae using the TRIzol reagent (Invitrogen, New York, NY, USA) according to the manufacturer’s instructions. The purity of RNA was quantified by a NanoDrop 2000 spectrophotometer (Thermo Fisher Scientific, New York, NY, USA). Poly (A)-tailed RNA prepared using magnetic oligo (dT) beads was broken into short fragments using a fragment buffer and was then reverse-transcribed to synthesize the first-strand cDNA with a random primer, and then DNA polymerase I was mixed with RNase H, dNTP, and the buffer solution to synthesize the complementary strand. The libraries were constructed using the Illumina methods and protocols, following the manufacturer’s instructions. The insert size and concentration of the cDNA library were both checked and quantified by an Agilent Bioanalyzer 2100 (Agilent Technologies, Inc., Santa Clara, CA) and Qubit^®^ RNA Assay Kit (Life Technologies, CA, USA), respectively. RNA-seq was carried out with an Illumina HiSeq 2500 instrument (Illumina, San Diego, CA, USA). In order to ensure the quality of information analysis, the raw reads were cleaned by removing the adapter sequences, empty reads, unknown nucleotides (ratio ≥10%), and low-quality reads with a basic mass value of Q ≤ 20 which accounted for more than 50% of the whole read length. The clean read assembly was performed according to the previous report [29]. Clean reads were mapped to the *B. mori* genome with the software package TopHat2 (version 2.0.12) [30]. The genome sequences and annotation file were downloaded from SilkDB. Then, the RNA-seq reads were aligned and the aligned reads were used to construct transcripts with Cufflinks (version 2.1.1) [31]. The functional annotation of differentially expressed gene (DEG) was performed using the gene ontology (GO) assignments [32], and Kyoto Encyclopedia of Genes and Genomes (KEGG) pathway enrichments [33].

### Tissue expression patterns of DEGs

The PGs are important endocrine organs that are significantly different from other tissues in both their morphology and function. The day-3 fifth instar of the silkworm is the boundary for the whole larval development stage [34, 35]. In order to analyze tissue expression patterns of the identified DEGs in PGs, we detected the expression patterns in multiple tissues of the head, integument, midgut, fat body, hemocyte, ovary, testis, malpighian tubule, trachea, ASG (anterior silk gland), MSG (median silk gland), and PSG (posterior silk gland) of day-3 fifth instar larvae. Total RNA was extracted using the TRIzol reagent (Invitrogen, Carlsbad, CA, USA). Total RNA concentrations were quantified and single-stranded cDNAs were synthesized. The *BmGAPDH* gene was used as an intrinsic control.

### Reverse transcription-quantitative PCR analysis

The genes selected according to the RNA-seq results were compared by reverse transcription-quantitative PCR (RT-qPCR). Total RNA was extracted from the PGs samples of the infected and control groups at 24, 48, and 72 h and the three developmental stages of day-3 (V3), day-6 (V6) and day-7 (V7) fifth instar larvae. The first strand cDNA was synthesized for use with the PrimeScript Reverse Transcriptase kit (TaKaRa, Dalian, China) according to the instructions of the manufacturer. RT-qPCR was performed as previously described [35]. The *BmGAPDH* gene was used as an intrinsic control.

## Results

### Statistics of precocious maturation of silkworm after infection with BmNPV

The proportion of mature silkworms being greater than 5% was considered to be the first gate and the majority of larvae that started maturing were considered as the second gate. Day-4 fifth instar larvae infected with BmNPV were matured early. The times that both the first gate and second gate mature silkworms appeared in the infected groups were 12 hours earlier than the control groups (**Fig 1**). The weights of mature silkworms in the infected groups were significantly decreased when compared with the control groups (**S2 Table**). The spinning process was normal and there was no difference between the infected groups and the control groups. About half of the larvae in the the infected groups died during the larval–pupal stage. Comparing with the control groups, the cocoon sizes and the weights of pupae (female and male) were observably reduced in the BmNPV-infected groups (**S2 Table** and **S3 Fig**). The fifth instar larvae underwent precocious maturation after infection with BmNPV. Moreover, the day-5 fifth instar larvae were fed with 8000 mg/L cholesterol and 7dC, respectively. The results of feeding with cholesterol and 7dC were also shown to induce precocious maturation when compared to the water control, where a certain number of larvae featured an anal prolapse in each group of feeding with cholesterol and 7dC (**S3 Table**).

**Fig. 1.**
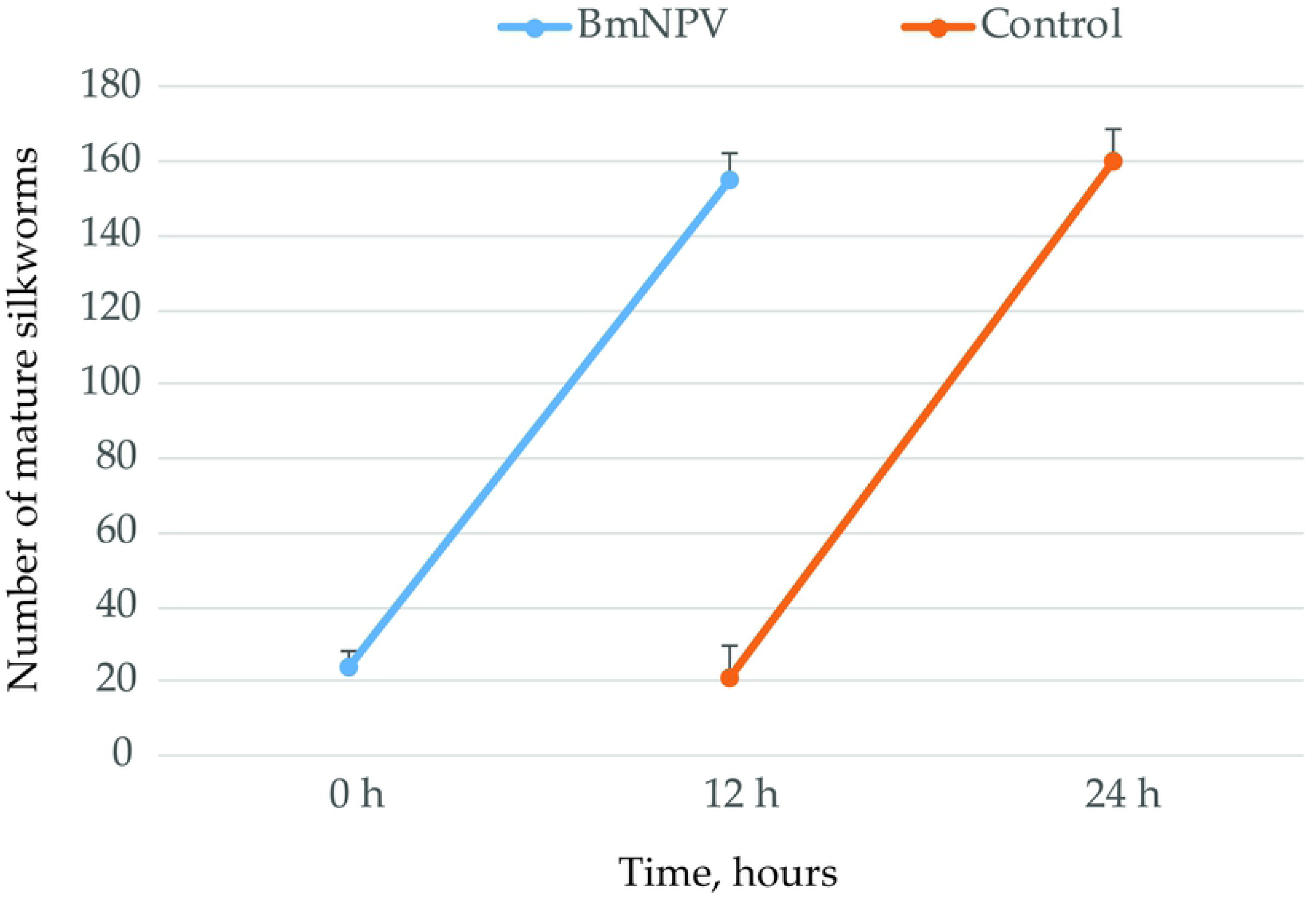
Statistics of precocious maturation for day-4 fifth instar silkworm larvae infected with *Bombyx mori* nucleopolyhedrovirus (BmNPV).

### Reverse transcription-PCR analysis of virus after BmNPV infection

Reverse transcription-PCR (RT-PCR) was used to analyze the virus genomic DNA copies in PGs at 24, 48, and 72 h after the BmNPV infection. The expressions of the *BmNPV IE1*, *GP41*, and *GP64* genes were detected in the PGs from 24 to 72 h after the virus infection (**Fig 2**). The expressions of the *IE1*, *GP41*, and *GP64* genes were not detected in PGs from the uninfected larvae at 48 h (**Fig 2**). The results reveal that the PGs were infected by the virus through oral inoculation. These results were useful for selecting the time point for the RNA-seq experiments.

**Fig. 2.**
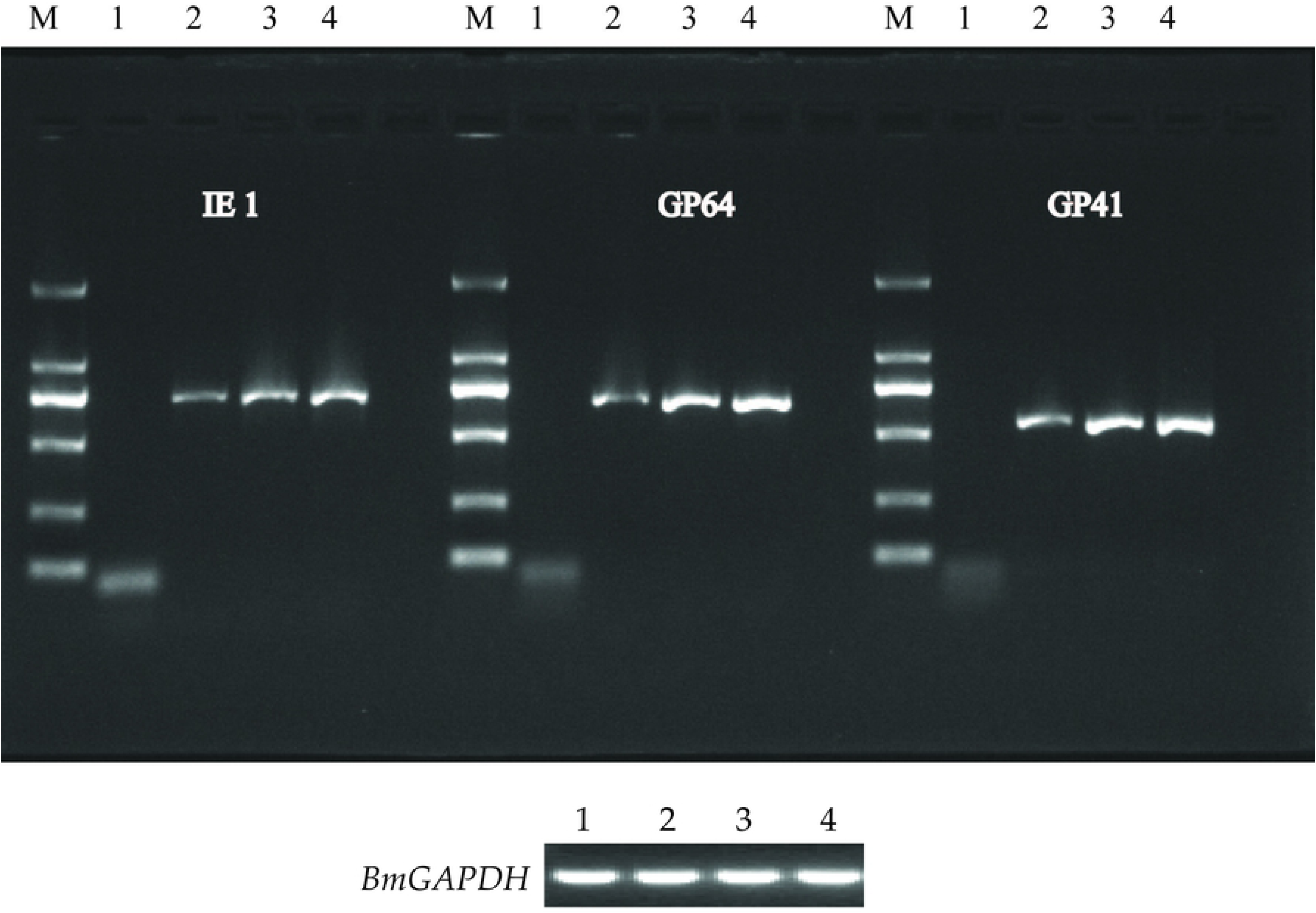
RT-PCR analysis of virus genomic DNA copies in prothoracic glands (PGs). The expressions of the BmNPV *IE1*, *GP41*, and *GP64* genes were detected in the PGs of the BmNPV-infected silkworm larvae at 24, 48, and 72 h, as well as the control larvae at 48 h. M: DL2000 DNA Maker; numbers 1 to 4 indicate control at 48 h and BmNPV-infected at 24 h, 48 h, and 72 h, respectively.

### Analysis of transcriptome data from the PGs

Transcriptome analysis is a widely used and effective technique to obtain a global overview of gene expression levels under different conditions. In this study, RNA-seq technology was used as a comparative transcriptomic approach to characterize some different features of PGs from the latter half of fifth instar larvae after BmNPV infection. The Pearson squared correlation coefficient (R^2^) between the four samples was greater than 0.938 (**S4 Fig**). The clean data of each group generated with a Q20 value were greater than 95% (**Table 1**). The paired-end clean reads were aligned to the *B. mori* genome with the TopHat2 (version 2.0.12) software package [30]. The percentage of clean sequences located on the genome was greater than 80% (**S4 Table**). These results indicate that the transcriptome data have been assembled with high quality and can be used for further research.

**Table 1.**
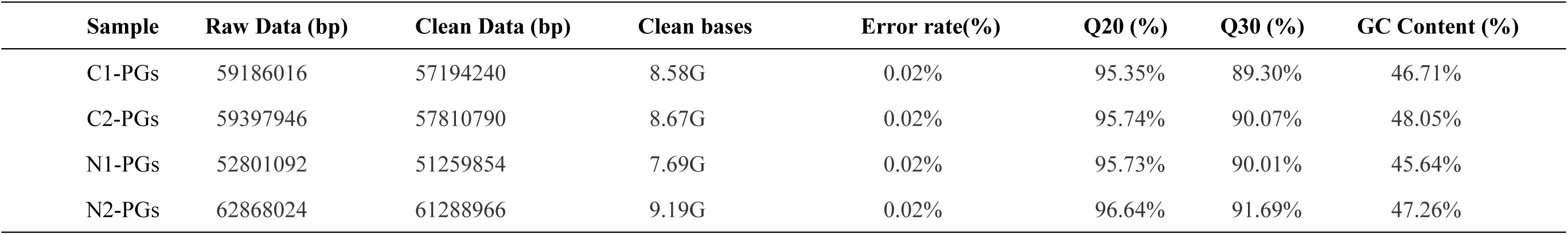
Summary statistics of prothoracic glands of *B. mori* genes based on transcriptome data

### Differentially expressed transcripts analysis and gene functional annotation

HTSeq (version 0.6.1) was used to count the reads numbers mapped to each gene for the quantification of gene expression levels [36]. Then, the fragments per kilobase of transcript per million mapped reads (FPKM) of each gene were calculated based on the length of the gene and read count mapped to a given gene. The clean reads were mapped to the *B*. *mori* genome. Genes with a FPKM ≥1.0 were identified as expressions. The number of expressed genes was 10,152 in the control groups and 10,404 in the BmNPV-infected groups (**S5 Table**). The up- and down-expressed transcripts were abstracted from the Cuffdiff results [31]. A ratio (log_2_ fold change) between the infected and control groups of ≥1.5 was identified as the determinant of differentially expressed genes (DEGs). In total, seven up-regulated and eight down-regulated DEGs were screened out (**Table 2**, **S6 Table**). The functions of the 15 DEGs were primarily located in the binding proteins of nucleic acids, ions, and proteins (**Table 2**). Then, the DEGs were annotated by gene ontology (GO) analysis to be involved in biological processes, molecular functions, and cellular components (**Fig 3**). The up-regulated expression genes were significantly enriched for genes related to biological processes that were basically focused on metabolic processes and biological processes (**Fig 3**, **S7 Table**). The down-regulated expression genes were basically focused on biological processes, metabolic processes, and in response to stress and stimulus in the biological processes of GO analysis (**Fig 3**, **S7 Table**). With regard to the cellular components, only the down-regulated expression genes were involved in the membrane and integral components of the membrane, and the up-regulated expression genes were not enriched (**Fig 3**, **S7 Table**). Within the molecular function, the up-regulated expression genes were primarily located in the catalytic activity and the down-regulated expression genes were involved in protein binding, hydrolase activity, and catalytic activity (**Fig 3**, **S7 Table**). There were some differences in the GO functional annotations between the up-regulated and down-regulated genes (**Fig 3**). The Kyoto Encyclopedia of Genes and Genomes (KEGG) pathway enrichment analysis of the identified DEGs showed that the enriched genes were mainly involved in pathways, including metabolic pathways, propanoate metabolism, beta-alanine metabolism, drug metabolism with other enzymes, pyrimidine metabolism, and pantothenate and coenzyme A (CoA) biosynthesis (**Table 3**, **S8 Table**).

**Fig. 3.**
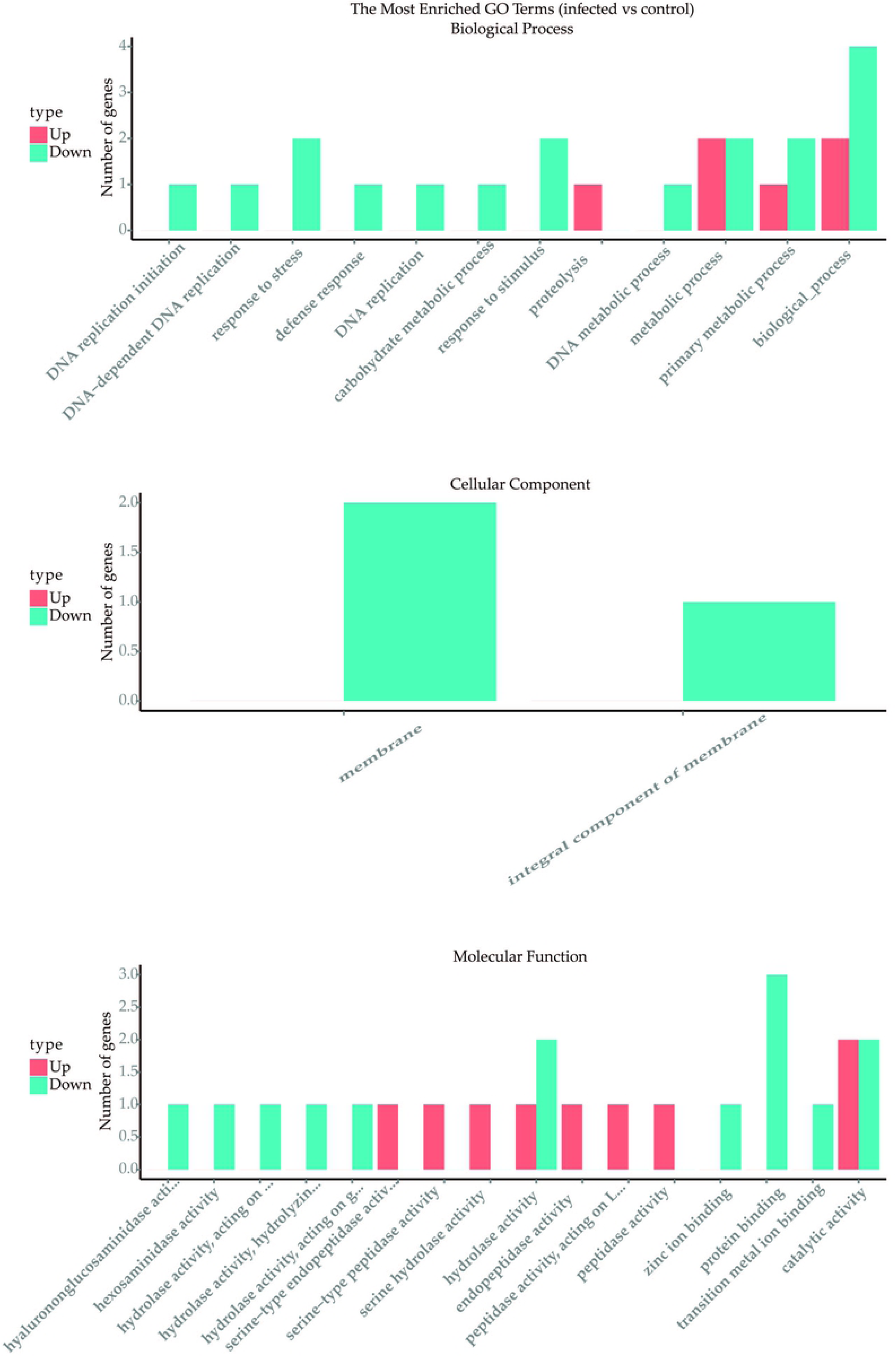
Gene ontology enrichment analysis for differential expression genes (DEGs). Genes are annotated by the biological process, cellular component, and molecular function.

**Table 2.**
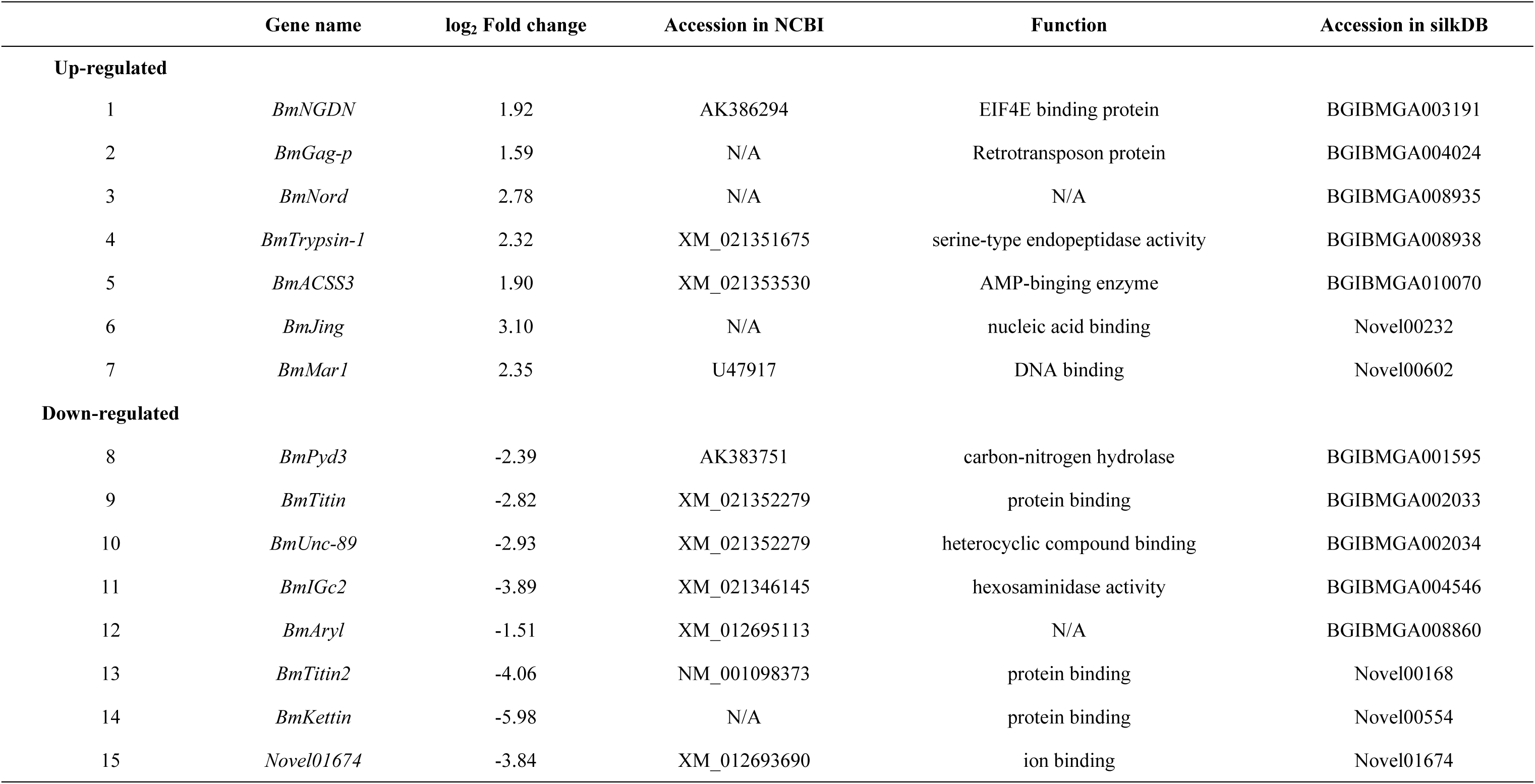
List of the differentially expressed genes in silkworm prothoracic glands with a 1.5-fold change after BmNPV infection

**Table 3.** The Kyoto Encyclopedia of Genes and Genomes (KEGG) pathway enrichment analysis

| Number | Map_Name | KEGG_ID | Gene_ID | Definition | log<sub>2</sub> Fold change |
| --- | --- | --- | --- | --- | --- |
| 1 | Metabolic pathways | bmor:100329149\| | BGIBMGA001595 | Beta-ureidopropionase [59] | -2.39 |
|  |  | bmor:101743386\| | BGIBMGA010070 | Fatty acid CoA ligase [60] | 1.90 |
| 2 | Propanoate metabolism | bmor:100329149\| | BGIBMGA001595 | Beta-alanine synthase [61] | -2.39 |
| 3 | Beta-Alanine metabolism | bmor:101743386\| | BGIBMGA010070 | Acyl-CoA synthetase [62] | 1.90 |
| 4 | Drug metabolism with other enzymes | bmor:100329149\| | BGIBMGA001595 | Cyanide hydratase [63] | -2.39 |
| 5 | Pyrimidine metabolism | bmor:100329149\| | BGIBMGA001595 | Beta-alanine synthase [61] | -2.39 |
| 6 | Pantothenate and CoA biosynthesis | bmor:100329149\| | BGIBMGA001595 | Pantetheine hydrolase [64] | -2.39 |

### Protein–protein interaction network analysis of BmACSS3

The KEGG pathway enrichment analysis of BmACSS3 (BGIBMGA010070) revealed acyl-coenzyme A (AcCoA) synthase activity. The orthologous groups of BmACSS3 in other species have the function of catalyzing the conversion of acetate into AcCoA, an essential intermediate at the junction of anabolic and catabolic pathways [37, 38]. BmACSS3 was submitted to the STRING database to generate protein–protein interaction networks. The main results of the enrichment network indicated that 10 proteins interact directly or indirectly with BmACSS3 (**Fig 4**, **S9 Table**). Functional enrichments in the network show 4 aldehyde dehydrogenase families (BGIBMGA010402-PA, BGIBMGA010403-PA, BGIBMGA001402-PA and BGIBMGA002457-PA) oxidizing a wide variety of aliphatic and aromatic aldehydes using nicotinamide-adenine dinucleotide phosphate (NADP) as a cofactor, namely, the 3 acyl-CoA acyltransferase 2 family (BGIBMGA012660-PA, BGIBMGA012661-PA and BGIBMGA014181-PA) with acetoacetyl-CoA and 3-ketoacyl-CoA thiolase activity, and the 3 AMP-binding enzyme family (BGIBMGA000721-PA, BGIBMGA008821-PA and BGIBMGA010070-PA) with catalyzing the conversion of acetate into AcCoA (**S8 Table**). The PGs are the main location of synthetic ecdysteroids from cholesterol and their subsequent secretion [5]. These results suggest that BmACSS3 may be involved in the acyl-CoA to cholesterogenesis pathways.

**Fig. 4.**
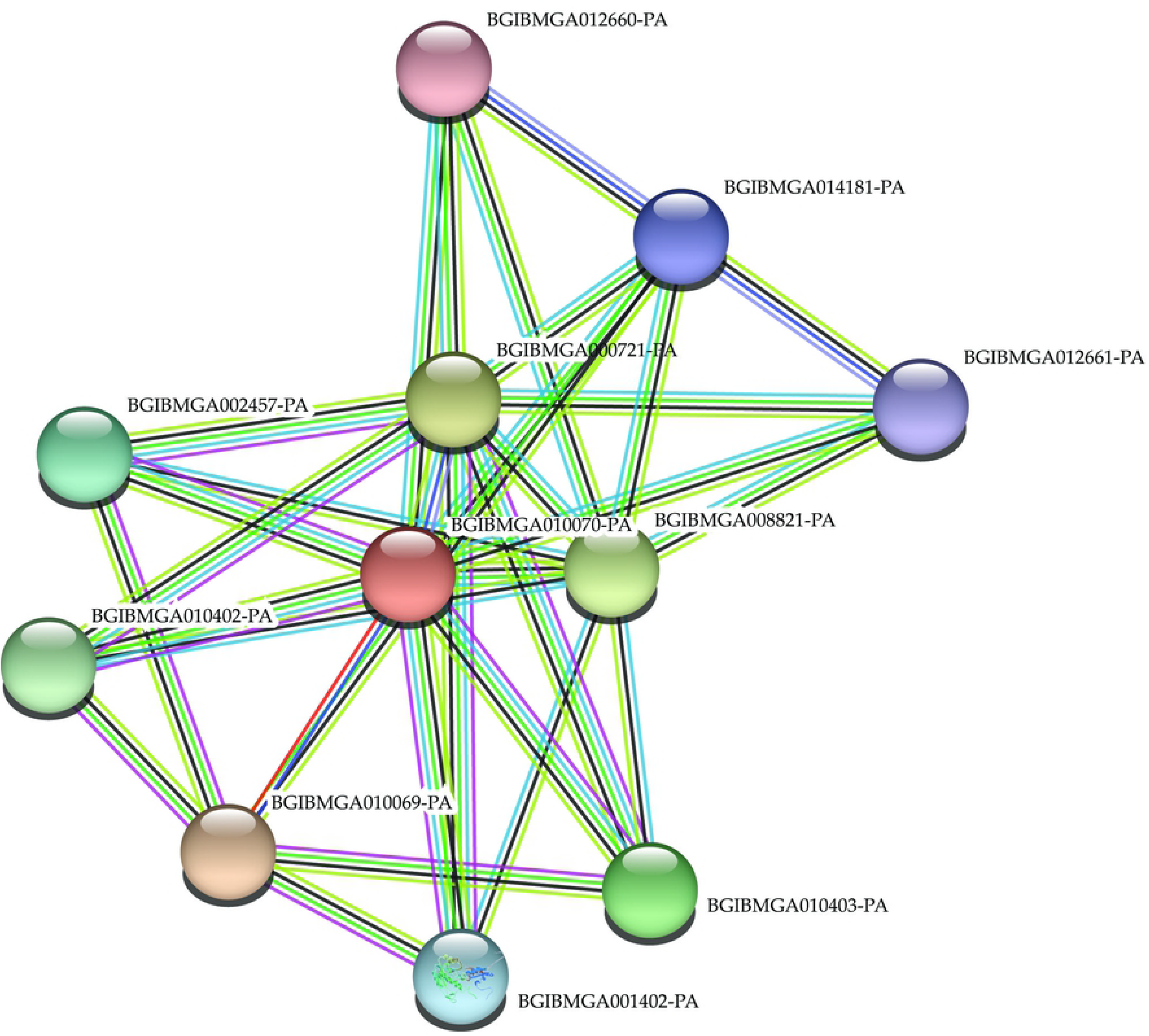
Protein–protein interaction networks of BmACSS3. Prediction and analysis of protein–protein interaction networks of BmACSS3 was supported by STRING.

### Spatial expression profile of the identified DEGs

The PGs are a model for synthetic ecdysone with regulating insect growth and development. The PGs are crowded with polygonal or oval-shaped secretory cells. The PGs are important endocrine organs that are significantly different from other tissues in both their morphology and function. The day-3 fifth instar of the silkworm is the boundary for the whole larval development stage [34, 35]. Thus, we investigated the spatial expression profile of the identified DEGs in PGs, including the gene expression levels across 12 *B*. *mori* larval tissues of day-3 fifth instars. The information of the DEGs and *BmGAPDH* primers is presented in **S1 Table**. The study will be helpful to understand the PGs and elucidate the expression characteristics of DEGs. The genes of *BmJing* and *BmAryl* had no expression signals in all 12 tissues (**Fig 5**). Novel01674 had a faint expression signal in malpighian tubules (**Fig 5**). *BmACSS3* was expressed only in the head and integument (**Fig 5**). The other 11 genes were expressed in multiple tissues (**Fig 5**).

**Fig. 5.**
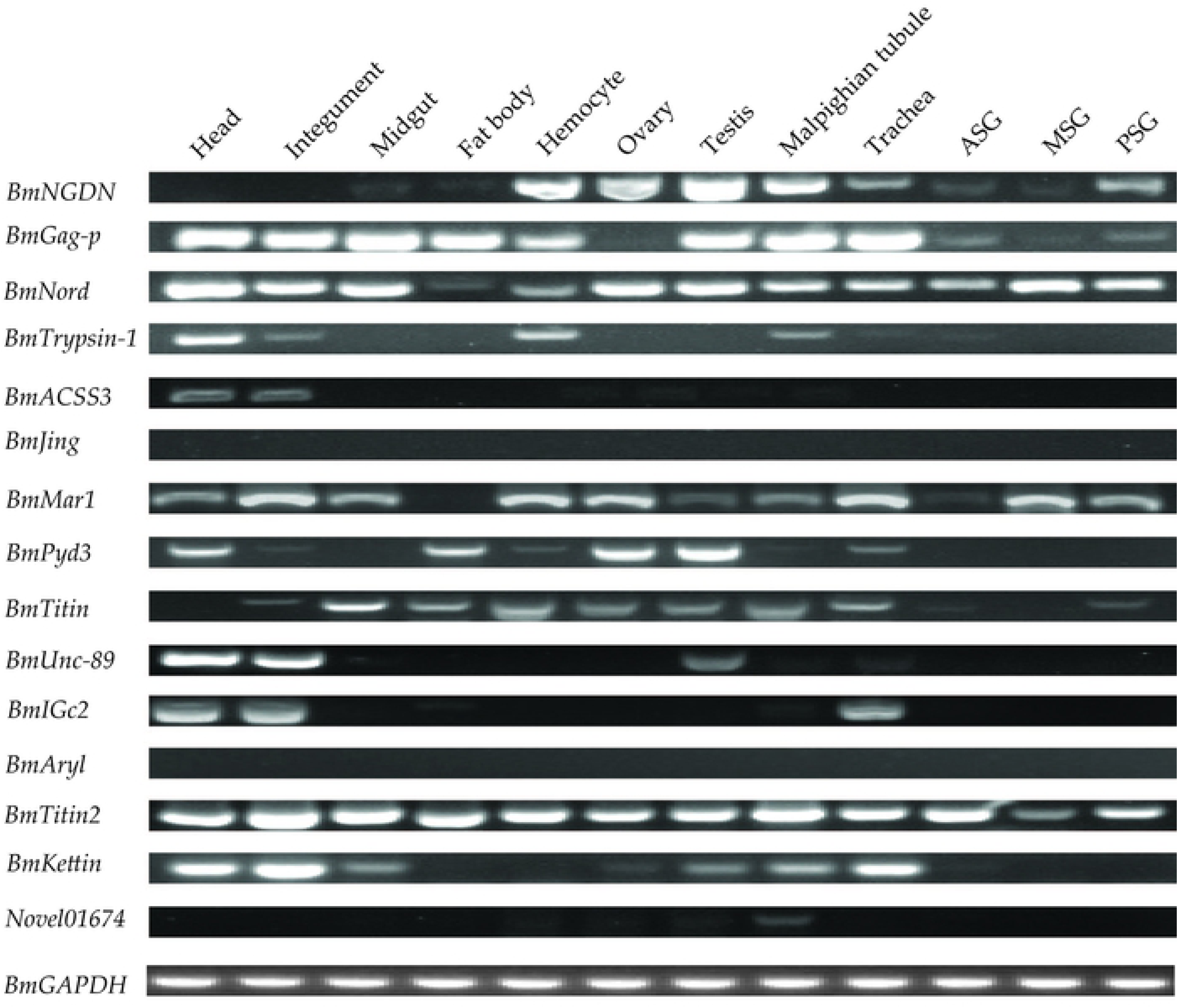
Expression patterns of DEGs in multiple tissues of day-3 fifth instar larvae. Reverse transcription (RT)-PCR was performed and the *BmGAPDH* gene was used as an internal control.

### Expression analysis of the identified DEGs

In the previous section, the day-4 fifth instar larvae infected by BmNPV matured early and the day-5 fifth instar larvae fed with cholesterol and 7dC also showed precocious maturation. Moreover, the day-3 fifth instar of the silkworm is the boundary for whole larval development stage [34, 35]. The ecdysteroid titer had a small rise and was followed by a plateau occurring one day before the silkworms started wandering. The titer started to increase gradually and elevated steeply to form a peak one day later on the next day (the majority of wandering) [39, 40]. According to the rearing conditions and the *B. mori* F_50_ strain, the silkworm larvae feed and grow quickly before the first half of the fifth instar, where the silkworm larvae massively synthesize silk proteins in the silk gland and get ready for the spinning and larval–pupal development throughout the latter half of the fifth instar. Day-7 (V7) fifth instar larvae of the *B. mori* F_50_ strain start the majority of the wandering. Therefore, we used reverse transcription-quantitative PCR (RT-qPCR) to investigate the relative expression levels of six randomly selected genes of the DEGs in the PGs of BmNPV-infected larvae at 24, 48, and 72 h and the three developmental stages of day-3 (V3), day-6 (V6), and day-7 (V7) for fifth instar larvae. A pair of specific primers for each gene was used in RT-qPCR, as shown in **S1 Table**, where the *BmGAPDH* gene was used as an intrinsic control. Here, six genes for three up- and down-regulated DEGs were selected, respectively. The expression levels of *BmNGDN*, *BmTrypsin-1*, and *BmACSS3* were up-regulated in the transcriptome data, their transcripts were significantly increased from 24 to 72 h after BmNPV infection (**Fig 6A–C**), and were also significantly increased in the developmental stages by V6 and V7 when compared to V3 (**Fig 6A1**, **B1**, **C1**). Meanwhile, the expression levels of *BmPyd3*, *BmTitin*, and *BmIGc2* were down-regulated in the transcriptome data, where their transcripts were significantly decreased from 24 to 72 h after BmNPV infection (**Fig 7A–C**) and were also significantly decreased in the developmental stages of V6 and V7 when comparing to V3 (**Fig 7A1**, **B1**, **C1**). In general, firstly, the changes in the six genes’ transcription were generally consistent with transcriptome data, and, secondly, they were all involved in the maturation process in the latter half of the fifth instar according to the characteristics shown in the above results, no matter whether the silkworm fifth instar larvae were infected by BmNPV or under the natural conditions.

**Fig. 6.**
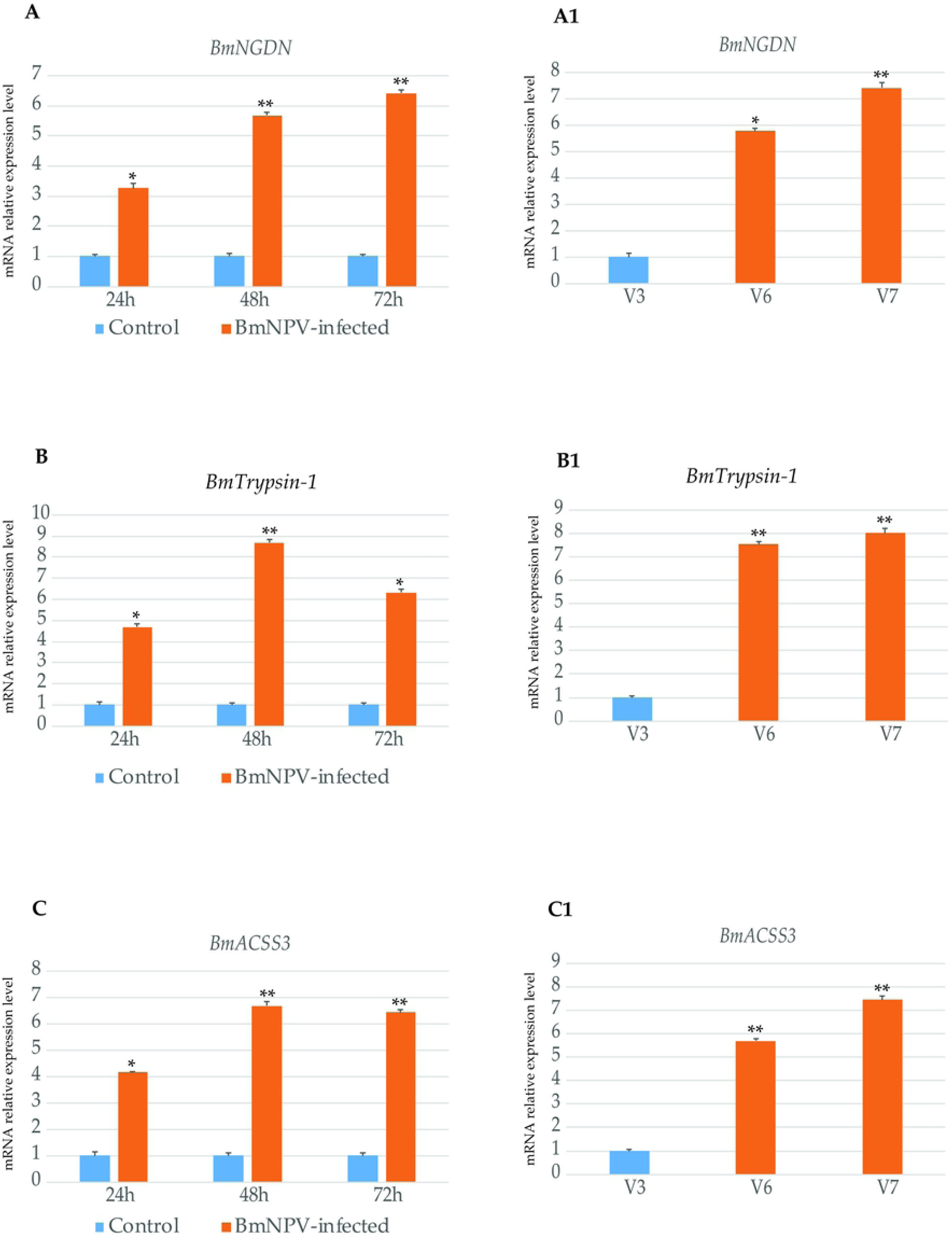
Analysis of the up-regulation genes from the transcriptome data of the **PGs of BmNPV-infected larvae and the developmental stages.** Here, *BmGAPDH* was used as internal control. The experiments were repeated three times. The data are the means ± SD of three independent experiments. The significant differences are indicated by * (p < 0.05) or ** (p < 0.01). Here, V3, V6, and V7, and day-3, day-6, and day-7 fifth instar larvae are featured.

**Fig. 7.**
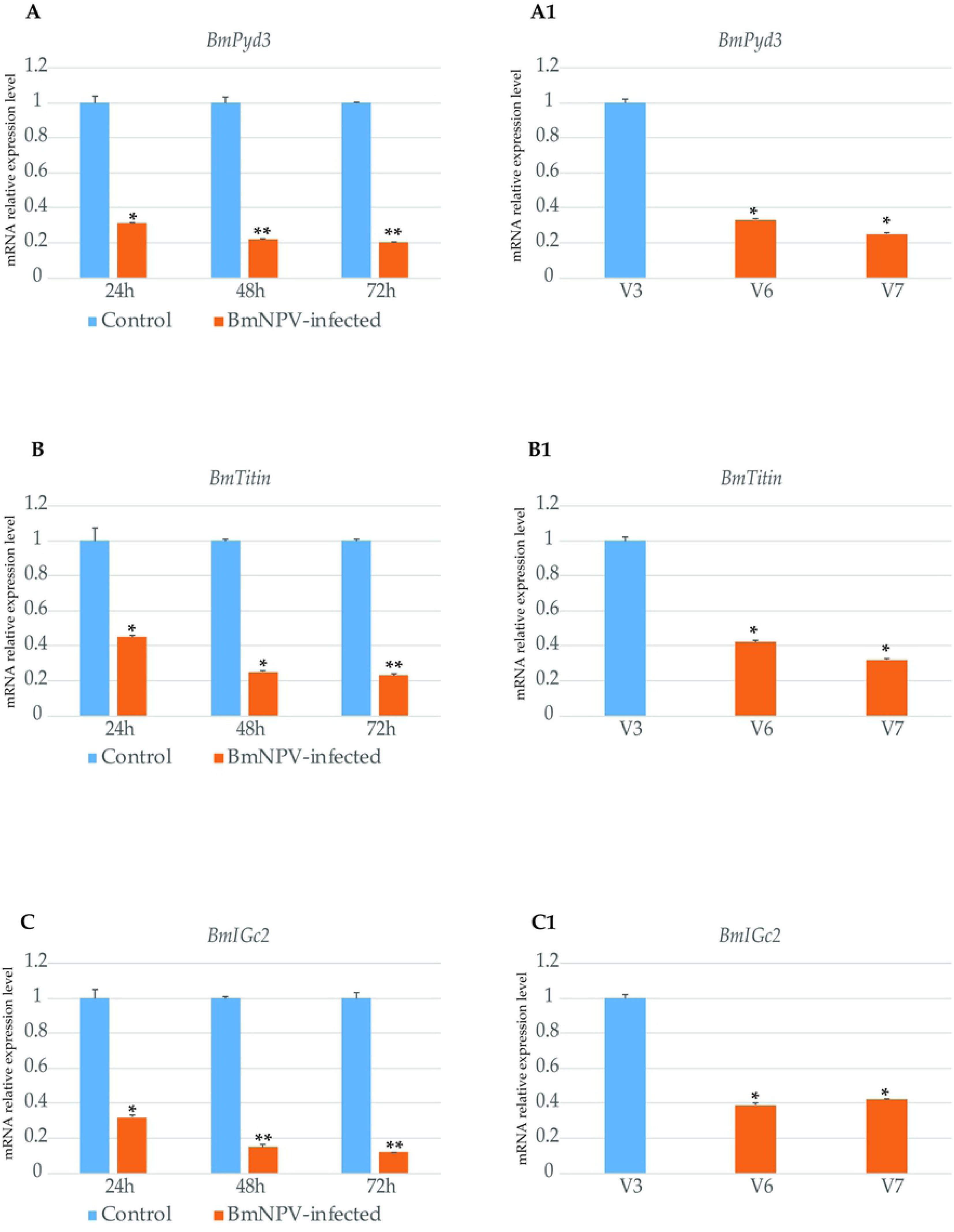
Analysis of the down-regulation genes from the transcriptome data of the PGs of BmNPV-infected larvae and the developmental stages. *BmGAPDH* was used as internal control. The experiments were repeated three times. The data are the means ± SD of three independent experiments. The significant differences are indicated by * (p < 0.05) or ** (p < 0.01). Here, V3, V6, and V7, and day-3, day-6, and day-7 fifth instar larvae are featured.

## Discussion

The silkworm, *B. mori*, is a complete metamorphosis insect, where one generation experiences full egg-larva-pupa-adult metamorphosis. In general, the larval stage needs to undergo four molting and five instars for a tetramolter silkworm. The last instar larva completes the larval–pupal transition. The developmental speed of the strain of the silkworm is regular and constant in the same rearing conditions. The silkworm is one of the best models to study insect physiology and biochemistry, especially to explore and study the relationship between induction factors (external and internal) and development.

In this study, day-4 fifth instar larvae infected by BmNPV matured early, and day-5 fifth instar larvae fed with cholesterol and 7dC also showed precocious maturation. The fifth instar period was shortened. In contrast, the fifth instar period was prolonged by feeding with cholesterol and 7dC in the first half of the fifth instar (day-1 to day-3 fifth instar), but no results were observed for BmNPV infection in the first half of the fifth instar because the larvae died. The nutrients needed for the vital activities were derived from the larval stage and this is the only time when the larva feeds during its whole lifecycle [35]. The amount of leaf ingested and digested quickly increases from immediately after ecdysis to the gluttonous stage in the first half of the fifth instar. Silkworm larvae greatly synthesize silk proteins in the silk gland and get ready for the spinning and larval–pupal development throughout the latter half of the fifth instar. The day-3 fifth instar (middle of this instar) of the silkworm is the boundary for whole larval development stage [34, 35]. Larvae mainly store essential nutrients for vital activities before the first half of this instar, and, in the latter half, larvae mainly synthesize massively silk proteins in the silk gland and get ready for larval–pupal development. Thus, we speculate that a reserve of basic nutrients for the fifth instar larvae is the prerequisite for developmental studies.

The chronic activation and ubiquitous expression of the innate immune pathogen recognition molecule of the peptidoglycan recognition protein (PGRP-LE) throughout the larval stages in *Drosophila* results in a severely reduced adult lifespan [41, 42]. In *Drosophila*, a strain of the obligate intracellular bacterium *Wolbachia pipientis* (*w*MelPop) has been described that shortens the life span of adult flies by up to 50%. *Drosophila* is the natural host of *Wolbachia* [43, 44]. The *Wolbachia* strain *w*MelPop has been successfully introduced into the mosquito *Aedes aegypti* and halves the mosquito adult lifespan [45]. The *w*MelPop infection induces the upregulation of the mosquito innate immune system and its presence inhibits the development of filarial nematodes in the mosquito [46]. The constitutive upregulation of immune genes is known to have a high cost and the cost of constitutive immune upregulation may contribute to the life-shortening phenotype [42, 46]. Moreover, insect innate immunity can be affected by juvenile hormones (JHs) and 20-hydroxyecdysone (20E) [47–49]. For instance, 20E induces antimicrobial peptides (AMP) and mRNA expression acting as an immune activator and JHs act as an immune suppressor by antagonizing 20E signaling in *Drosophila* [47, 48, 50, 51]. In contrast to *Drosophila*, JHs act as an immune activator, while 20E inhibits innate immunity in the fat body during *B. mori* postembryonic development [49]. Molting and metamorphosis are coordinately regulated by the molting hormone 20E and JHs in insects. Briefly, the prothoracic glands (PGs) release ecdysteroids that induce larval or metamorphic ecdysis, depending on the presence of the JH secreted by the corpora allata [40]. The PGs are important endocrine organs and known to be cholesterol-rich tissue [1]. They are known as the main site of synthetic ecdysteroids derived from cholesterol and their subsequent secretion [5, 6].

In this study, the RT-PCR results confirmed that the silkworm PGs were infected by BmNPV through oral inoculation. Then, the RNA of silkworm PGs was extracted and sequenced by RNA-seq 48 hours after BmNPV infection. There were seven up-regulated and eight down-regulated DEGs that were identified. The classifications of the 15 DEGs were primarily located in binding activity of nucleic acids, ions, and proteins which were mainly involved in the metabolic processes and pathways. The spatial expression profiles of *BmJing* and *BmAryl* had no expression signals in all 12 tissues. The genes of *BmJing* and *BmAryl* may be specifically expressed in the silkworm PGs. The RT-qPCR results of the DEGs in the PGs of BmNPV-infected at 24, 48, and 72 h were generally consistent with the transcriptome data, which may be related to the BmNPV infection promoting the early maturation in the latter half of the silkworm fifth instar. The ecdysteroid titer had a small rise and was followed by a plateau occurring one day before the silkworms started wandering, where the titer started to increase gradually and elevated steeply to form a peak one day later on the next day (the majority of wandering) [39, 40]. The expression analysis of the DEGs in the PGs of the fifth instar larvae developmental stages by day-6 (V6) and day-7 (V7), comparing to day-3 (V3), was basically the same as that of the BmNPV infection, promoting the early maturation in the latter half of the silkworm fifth instar. The identified DEGs might be involved in the maturation process in the latter half of the fifth instar.

Steroid hormones play a pivotal role in many biological processes in insects, especially in guiding the transition from one developmental stage to the next via molting and metamorphosis. PGs are known to be cholesterol-rich tissue and are the main site of synthetic ecdysone from dietary cholesterol or phytosterols via a series of hydroxylation and oxidation steps [5, 6]. Fatty acids are the building blocks of many lipids, including triacylglycerol and cholesteryl esters [52]. For this, acyl-coenzyme A synthetases (ACSs) catalyze fatty acids to form a thioester with CoA, which is a common initial step of all fatty acid metabolic processes [53]. The acyl-CoA is the primary cholesterogenesis substrate [38]. In *Drosophila*, pudgy is an acyl-CoA synthetase associated with mitochondria that activates fatty acids for beta-oxidation [54] and RNAi-mediated knockdown of the acyl-CoA synthetase gene, also prolonging and extending maximum lifespan [55]. Moreover, the trypsin modulating oostatic factor (TMOF) has the ability to inhibit trypsin biosynthesis [56], TMOF is able to inhibit ecdysone biosynthesis in the larval ring glands and the PGs, causing a delay in pupariation [57, 58]. In silkworm PGs, the transcripts of *BmTrypsin-1* and *BmACSS3* were significantly increased from 24 to 72 h after BmNPV infection and in the developmental stages by V6 and V7 when compared to V3. *BmTrypsin-1* and *BmACSS3* may be involved in the maturation process in the latter half of silkworm fifth instar larvae.

## Conclusion

The silkworm is a complete metamorphosis insect, where one generation features full egg-larva-pupa-adult metamorphosis, which makes the insect a suitable model to study insect physiology and biochemistry. BmNPV is a principal pathogen of the silkworm and its host range is restricted to silkworm larvae. In this study, BmNPV-infected day-4 fifth instar larvae matured early, and day-5 fifth instar larvae were fed with cholesterol and 7dC and also showed precocious maturation. The PGs are a model for synthetic ecdysone for regulating insect growth and development. We used RNA-seq to analyze silkworm PGs after BmNPV infection, where seven up-regulated and eight down-regulated DEGs were identified. The classifications of the 15 DEGs were primarily involved in the metabolic processes and pathways. The spatial expression profiles of *BmJing* and *BmAryl* indicated that they may be specifically expressed in the silkworm PGs. The RT-qPCR results of the DEGs in the PGs of BmNPV-infected larvae after 24, 48, and 72 h and during the developmental stages by day-6 (V6) and day-7 (V7), comparing to day-3 (V3), revealed that the DEGs may be related to the BmNPV infection, which promotes early maturation in the latter half of the silkworm fifth instar. To our knowledge, this study is the first report on the identification of possible genes in PGs correlating with the precocious molting and metamorphosis of silkworm larvae under BmNPV infection in the latter half of the fifth instar. Our findings expand the current knowledge of the complex biological processes in the interactions between BmNPV and host precocious metamorphosis. This work provides a new perspective on BmNPV infection and host developmental response, as well as suggesting candidate genes for further research.

## Supporting information

**S1 Figure View of the PGs entwined in the tracheal bush of the first spiracle.**

**S2 Figure View of the PGs from the fifth instar silkworm larva.**

**S3 Figure Effects of BmNPV-infected in the weight of mature silkworms and pupae.** (A) Dorsal view of the female pupae that were infected with BmNPV to compare with the control. (B) Dorsal view of the male pupae that were infected with BmNPV to compare with the control. (C) View of the cocoons that were infected with BmNPV to compare with the control.

**S4 Figure Analysis of the correlation of RNA-seq data.** (A) Diagram of correlation coefficient between samples. (B) Correlation between the C1 and C2. (C) Correlation between the C1 and N1. (D) Correlation between the C1 and N2. (E) Correlation between the C2 and N1. (F) Correlation between the C2 and N2. (G) Correlation between the N1 and N2. C1 and C2 indicate two independent biological experiments of transcriptome sequencing of PGs in the control groups, respectively. N1 and N2 indicate two independent biological experiments of transcriptome sequencing of PGs in the control groups, respectively.

**S1 Table Primers of genes for RT-PCR and RT-qPCR**

**S2 Table The weight of mature silkworms and pupae in BmNPV-infected and control groups**

**S3 Table The results of the feeding experiment with cholesterol, and 7-dehydrocholesterol (7dC), using water as control**

**S4 Table A list of reads compared to the *Bombyx mori* genome**

**S5 Table A list of genes with FPKM ≥1.0 between infected and control groups**

**S6 Table A list of differential expression genes between infected and control groups**

**S7 Table Infected vs. control DEG_GO_enrichment result**

**S8 Table Infected vs. control all DEG_KEGG_pathway_enrichment result**

**S9 Table Protein–protein interaction annotation**

## Acknowledgments

This research was financially supported by the Jiangsu Provincial Natural Science Foundation of China (grant no. BK2012273) and the National Natural Science Foundation of China (grant no. 31302035, 2735011802).

## Author contributions

Pingzhen Xu performed the experiment and wrote the manuscript. Meirong Zhang performed the literature review and analyzed the data. Xueyang Wang prepared the illustrations and collected the data; Yangchun Wu suggested important research points. All authors have read and approved the final version of the manuscript. The authors declare that they have no conflicts of interest related to this publication.

